# An Approximate Markov Model for the Wright-Fisher Diffusion

**DOI:** 10.1101/030940

**Authors:** Anna Ferrer-Admetlla, Christoph Leuenberger, Jeffrey D. Jensen, Daniel Wegmann

## Abstract

The joint and accurate inference of selection and demography from genetic data is considered a particularly challenging question in population genetics, since both process may lead to very similar patterns of genetic diversity. However, additional information for disentangling these effects may be obtained by observing changes in allele frequencies over multiple time points. Such data is common in experimental evolution studies, as well as in the comparison of ancient and contemporary samples. Leveraging this information, however, has been computationally challenging, particularly when considering multi-locus data sets. To overcome these issues, we introduce a novel, discrete approximation for diffusion processes, termed *mean transition time approximation*, which preserves the long-term behavior of the underlying continuous diffusion process. We then derive this approximation for the particular case of inferring selection and demography from time series data under the classic Wright-Fisher model and demonstrate that our approximation is well suited to describe allele trajectories through time, even when only a few states are used. We then develop a Bayesian inference approach to jointly infer the population size and locus-specific selection coefficients with high accuracy, and further extend this model to also infer the rates of sequencing errors and mutations. We finally apply our approach to recent experimental data on the evolution of drug resistance in Influenza virus, identifying likely targets of selection and finding evidence for much larger viral population sizes than previously reported.

---

Detecting signatures of past selective events provides insights into the evolutionary history of a species and elucidates the interaction between genotype and phenotype, offering important functional information. Unfortunately, a population’s demographic history is a major confounding factor when inferring past selective events, particularly because demographic events can mimic many of the molecular signatures of selection (Andolfatto and Przeworski 2000; Nielsen 2005). Despite efforts to create statistics robust to demography, all currently available methods to detect selection are prone to mis-inference under non-equilibrium demography.

Some of these issues can potentially be overcome by using multi-time point data, as the trajectory of even a single allele contains valuable information about the underlying selection coefficient. Owing to advances in sequencing technologies, such multi-time point data are becoming increasingly common from experimental evolution (Foll *et al*. 2014a), from longitudinal medical or ecological studies (Wei *et al*. 1995; Renzette *et al*. 2013), and through ancient samples (Wilde *et al*. 2014; Sverrisdóttir *et al*. 2014). However, computationally efficient and accurate methods to infer demography and selection jointly from such data sets are still limited.

A natural and common way of modeling such time series data is in a Hidden Markov-Model (HMM) framework, which allows efficient integration over the distribution of unobserved states of the true population frequencies, thus allowing calculation of the likelihood based on the observed samples. Williamson & Slatkin (1999) (Williamson and Slatkin 1999), for instance, developed a maximum-likelihood approach based on such an HMM to infer the population size *N* from samples taken at different time points. More recently, similar approaches have been developed to infer population size along with the selection coefficient of a selected locus for which time series data is available (Bollback *et al*. 2008; Malaspinas *et al*. 2012).

All such approaches, however, are plagued by the problem that the number of hidden frequency states is equal to the population size, which renders HMM applications computationally unfeasible for large populations. Different routes have been taken to overcome this. One approach is to model the underlying Wright-Fisher process as a continuous diffusion process, which is then discretized for numerical integration using a numerical difference scheme (Bollback *et al*. 2008). Since, this approach remains computationally expensive, it was later suggested to directly model the diffusion process on a more coarse-grained grid (Malaspinas *et al*. 2012). Under this approach, their generator matrix for the transition between the coarse grained states is then approximated by fitting the first and second infinitesimal moments. Unfortunately, the minimum number of states required is still computationally prohibitive for large values of *γ* = 2*Ns* (Malaspinas *et al*. 2012). For this reason, the most recent reported method resorted to simulation based Approximate Bayesian Computation (ABC), which allowed the joint inference of locus-specific selection coefficients for many loci (Foll *et al*. 2014a,b). However, this method requires first estimating the population size under the assumption that all loci are neutral, and thus may be biased when many loci are under selection.

Here we introduce a novel framework by approximating the WF-process with a coarse-grained Markov-Model that exactly preserves the expected waiting times for transition between states. This is achieved by exploiting the theory of Green’s function for diffusion processes. Contrary to previous approaches, our approximation matches the WF-process closely even when only very few states are considered, regardless of *γ =* 2*Ns*. As we show with extensive simulations and a data application from experimental evolution, our method allows for accurate joint inference of both population size and locus-specific selection coefficients even in the presence of pervasive selection. Further, it is readily extended to incorporate population size changes, sequencing errors or the appearance of novel mutations.

## Models

### Mean Transition Time Approximation

Let *X*(*t*) be a diffusion process on the state space [0,1]. This is a continuous-time Markov process with continuous sample paths and with infinitesimal generator

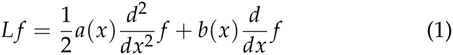

The classical example in population genetics is the Fisher-Wright diffusion which we discuss below. We seek to find a discrete-state Markov process *U*(*t*) which approximates *X*(*t*). For this purpose, we subdivide the unit interval [0,1] into, not necessarily equidistant, frequencies

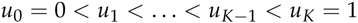

These form the states of *U*(*t*). For two states *u_i_*, *u_j_*, consider the transition time to first visit of *u_j_* when starting at *u_i_*.

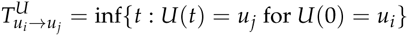

Similarly we define the transition time for the diffusion process *X*(*t*). We say that *U*(*t*) is a *mean transition time approximation* of *X*(*t*) if

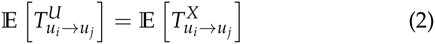
for all pairs of states *u_i_*, *u_j_* (see Fig. 1). This condition guarantees that the paths of *X*(*t*) and *U*(*t*) exhibit comparable long-time behavior. In the following we show how to construct the Markov process *U*(*t*) from the diffusion process *X*(*t*) using the theory of Green’s function.

**Figure 1.**
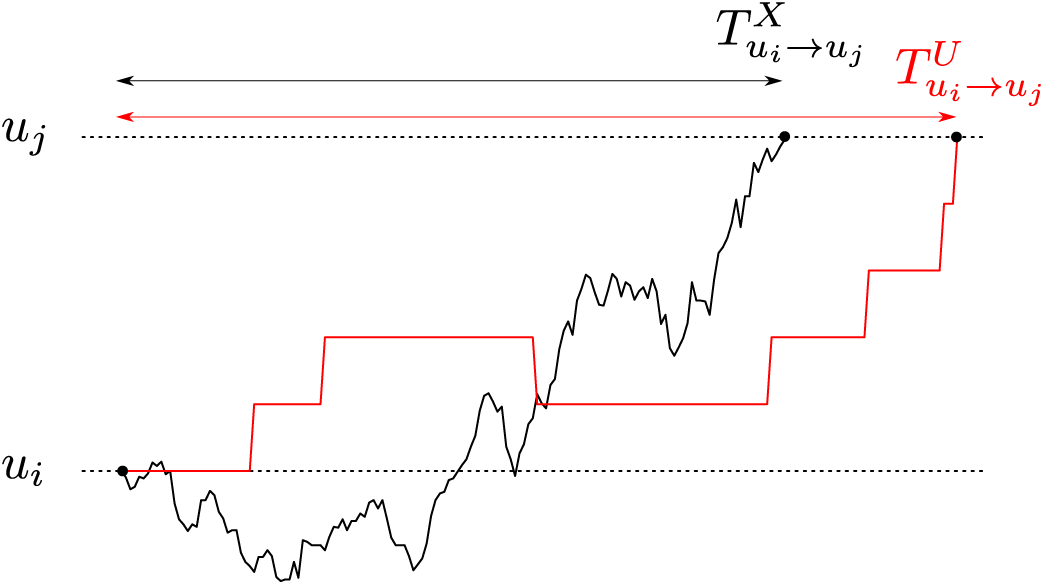
Mean transition time approximation of Markov processes. Shown are the realizations of a continuous diffusion process *X*(*t*) (black) and a discrete-state Markov process *U*(*t*) (red) starting at *u_i_* until they reach *u_j_* for the first time. If the expected waiting time for such a transition is the same for both process for all pairs of states *u_i_, u_j_*, we say that *U*(*t*) is a mean transition time approximation of *X*(*t*).

We begin by recalling some notions for diffusion processes. The natural scale of the process *X*(*t*) is given by

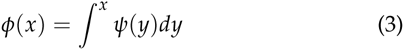
where 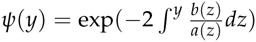. The so-called speed measure is defined by

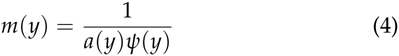

The Green’s function for an interval (*u, v*) ⊆ [0,1] is given by

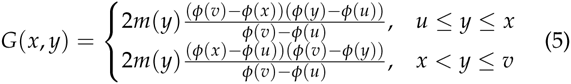

Denote by T*_x→u_* or T*_x→v_* the time to first visit of *u* or *v*, respectively, starting at *x*. Then 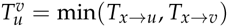 is the exit time from the interval (*u*_1_, *u*_2_), given the process is at *x* at time *t =* 0. One can show

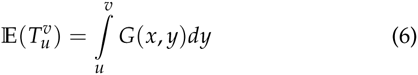

Moreover, the probability of exiting at the lower limit *u* is

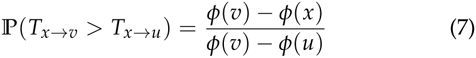

We now want to determine the instantaneous transition rates *q_i,j_* of the discrete-state Markov process *U*(*t*). Recall the definitions

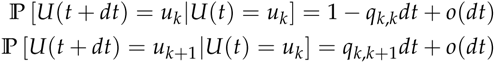
and

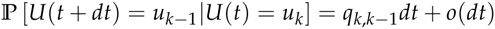

The sojourn time of state *u_k_*, i.e. the time interval of *U*(*t*) spent in state *u_k_*, is an exponential random variable with parameter *q_k,k_*. Since the expectation of this exponential variable is 1/*qk,k* our condition (2) enforces

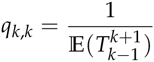
where we write *k* − 1 instead of *u_k_* etc. in order to unburden the notation. From this we get

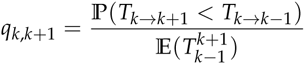
and

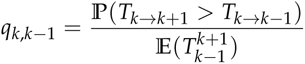

We can now form the tridiagonal generator matrix

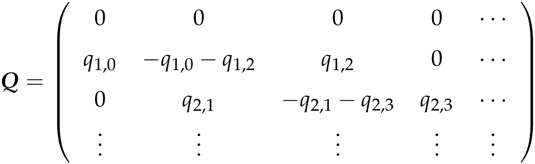

The transition matrix of the Markov process *U*(*t*) is given by

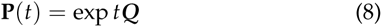

### Application to Wright-Fisher Models

We will consider a classic Wright-Fisher Model of two alleles *A* and *a* with fitnesses *s* and 1 − *s*, respectively, that segregate in a population of size 2*N*. Time *t* is measured in generations of the Wright-Fisher process. In the presence of a non-vanishing dominance coefficient *h* the fitnesses of the three genotypes are given by *w_AA_* = 1 + *s*, *w_Aa_* = 1 + *hs*, and *w_aa_* = 1. Under such a model, the infinitesimal mean, which corresponds to the change in allele frequency, is then given by

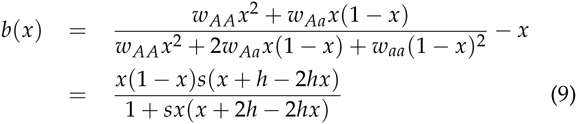

Let *X*(*t*) be a diffusion process corresponding to the frequency of allele *A*. An excellent approximation of the Wright-Fisher process is obtained by setting

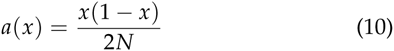
and

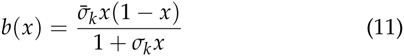
in the infinitesimal generator (1), where

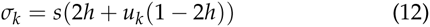
and

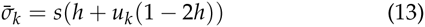
when *u_k−_*_1_ ≤ *x ≤ u_k+_*_1_

Note that in the standard diffusion approximation the denominator term in *b*(*x*) is often omitted. But as shown in (Lacerda and Seoighe 2014), the above choice yields a much more accurate approximation to the WF process.

From (3) and (4) we get

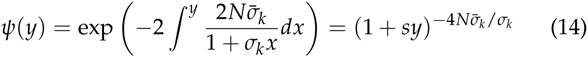
and

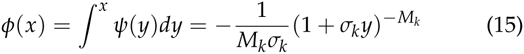
where we have set

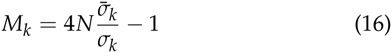

For the speed measure we obtain

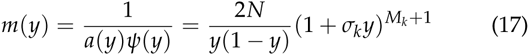

Consider three consecutive states *u_k−_*_1_, *u_k_* and *u_k+_*_1_. For the probability to exit at the lower state we get

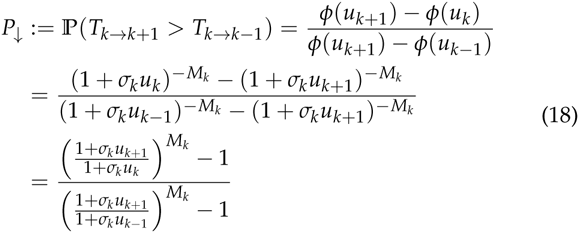

The probability for exit at the upper state is

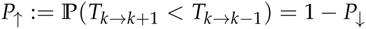

Observe that the Green’s function is calculated by

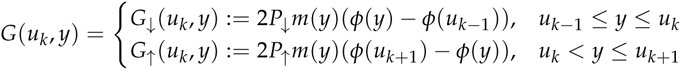

Using the quantities calculated above we get for the two parts of the Green’s function:

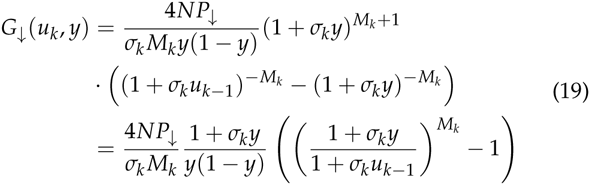
and

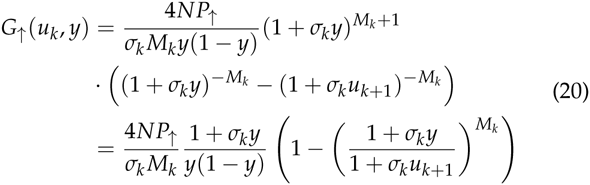

With numerical integration we can determine

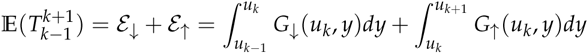

Specifically, we use the extended Simpson’s rule for the numerical integration (Press 2007), which we found to give accurate results with typically only 8 or 10 intervals.

If *γ =* 2*N*s is large, we get approximations for the Green function which allow for analytic expressions of the integrals (see Appendix). Similarly, analytical expressions can be found in the special case *s* = 0 (see Appendix).

### Bayesian Inference

Consider that at the times *Tt, t* = 0,…, *T*, samples of sizes *M_t_* were taken from the population and *m_t_* alleles A were observed in these samples. In this section, we describe how the mean transition time approximation introduced above can be embedded into a Bayesian inference scheme to estimate the population size 2*N* and the locus-specific selection coefficient jointly from time series data.

As has been noted previously (Williamson and Slatkin 1999; Malaspinas *et al*. 2012; Mathieson and McVean 2013; Lacerda and Seoighe 2014), a natural way of modeling both the underlying evolutionary process as well as the process of sampling is a Hidden Markov Model (HMM). Under the assumption that the population size between two time points *T_t_* and *T_t_*_+1_ is constant at *N_t_*, the transition matrix of such an HMM from state *U*(*T_t_*) to state *U*(*T_t_*+1) is calculated by

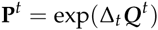
where Δ*_t_* = *T_t_*_+1_ − *T_t_* and the generator matrix *Q^t^* is determined as explained above using *N = Nt*. We note here that this framework allows for instantaneous population size changes to occur at every time *t* during the HMM. However, we will henceforth only deal with situations in which the population size is assumed to be constant across the whole sampling period.

Following previous implementations (e.g. (Williamson and Slatkin 1999; Malaspinas *et al*. 2012; Mathieson and McVean 2013; Lacerda and Seoighe 2014)), we will assume that the sampling of alleles from the underlying population frequency is binomial, i.e.

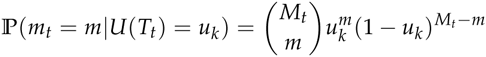

However, for large sample sizes, the few states *u_k_* may be too coarse grained to capture the region of high emission probability. We thus propose to integrate the emission probabilities against a smoothing kernel. we chose to implement a beta distribution kernel, which leads to a beta-binomial emission probability that can be evaluated analytically. Specifically, we chose to use a beta-kernel with mean *u_k_* and variance 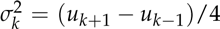. Under this choice, the emission probabilities are then calculated by

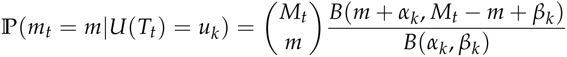
where B(·,·) is the Beta function and the parameters *α_k_* and *β_k_* are determined via the moment estimators for a beta distribution

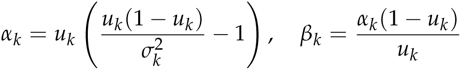

With both transition and emission probability matrices at hand, we calculate the likelihood of the full data using the standard forward recursion. To be specific, let us first define for *t* = 0,…, *T* the (*K* + 1) × (*M_t_* + 1) emission probability matrices

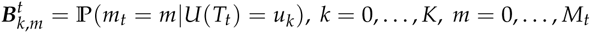

Denoting ***m***_1_*_:t_* = (*m*_1_,…, *m_t_*), we define the total probability

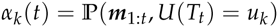

This total probability can be determined efficiently with the forward recursion

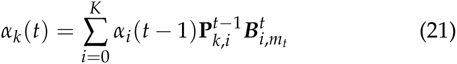
and 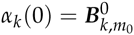. Then one has

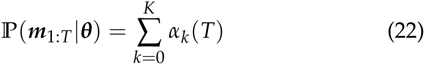
where we made explicit the dependence of this probability on the parameters

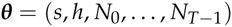
If we impose priors *π*(***θ***) on the parameters then we can simulate the posterior probability *π*(**θ**|***m*_1:_*_T_***) with the usual MCMC scheme using (22) and the Hastings ratio

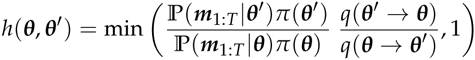

### Extension of basic model

**Sequencing errors** Generally, sequencing errors are overcome with sufficient coverage. However, in many applications of next generation sequencing to experimental evolution, the goal of the sequencing is not to infer individual genotypes, but rather allele frequencies directly. Under such a setting, each sequencing read is assumed to be from a different individual. In such cases, sequencing errors may lead to false inference, especially when allele frequencies are very small.

Incorporating sequencing errors into our framework is straight forward. Under the assumption that there are only two alleles present at the locus (achieved by, for instance, pooling all non-selected alleles into one class) and symmetric error rates *ϵ* between those classes, we can approximate the probability that 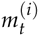, the *i*-th allele survey at time *t*, is A in the presence of sequencing errors as

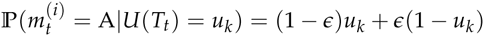

**Mutational Input** We allow for the production of mutant alleles only when the process is in state *u*_0_ = 0 or *u_K_* = 1. The production of new alleles proceeds at a rate of 2*Nµdt*. Once a new allele is produced, say when the system is in state *u*_0_, it must get from state 1 / 2*N* into state *u*_1_. This happens with probability 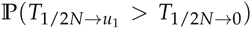 which is calculated according to (7). This yields the transition rate

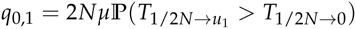

In the case of positive selection we have

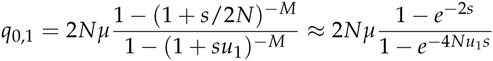

With *M =* 4*N −* 1. A similar logic yields

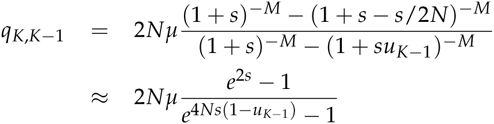

In the selection-free case the transition probabilities are much simpler:

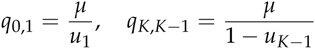

### Implementation

We have implemented the proposed model and the Bayesian inference scheme in an easy-to-use C++ program available on our lab website. While we use standard implementations for most aspects, we note the matrix exponentiation in Eq. 8, which is a numerically very demanding problem. A classic algorithm for matrix exponentiation is by diagonalization of the matrix (Moler and Van Loan 1978). While computationally efficient, this algorithm may be numerically unstable for matrices with large conditions numbers, which are typically observed when *γ =* 2*Ns* becomes large. This was previously observed by (Malaspinas *et al*. 2012), who addressed this issue using multiple precision arithmetics. Unfortunately, such arithmetics are computationally very demanding, leading to slow performance of their implementation.

Here, we propose to alleviate this problem using the approximation

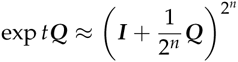
which can be calculated by successive quadration. Such matrix multiplications are generally demanding, but can be implemented in a computationally efficient manner for generator matrices that are tridiagonal, as each quadration step only adds two additional diagonals and such band matrices can be multiplied efficiently (see chapter 7.4 in (Dahlquist and Björk 2008)).

### Application to Influenza data

We analyzed allele frequency data from whole genome data sets of Influenza H1N1 obtained in a a recent evolutionary experiment (Foll *et al*. 2014a). While we refer the reader to the original study for a detailed description of the experimental set-up, we summarize the key point briefly here: Influenza A/Brisbane/59/2007 (H1N1) was serially amplified on Madin-Darby canine kidney (MDCK) cells for 12 passages of 72 hours each to prevent any freeze-thaw cycles. After the three initial cycles, samples were passed either in the absence of drug, or in the presence of increasing concentrations of oseltamivir, a neuraminidase inhibitor for another 9 passages. At the end of each passage, samples were collected for whole genome high throughput population sequencing up to a median coverage of more than 50,000x.

For our analysis here we only considered the time-points taken during drug treatment (passages 4 to 12). As in (Foll *et al*. 2014a), we excluded all monomorphic sites and those with a coverage lower than 100, resulting in a total of 1427 included sites. For each site, we only kept the two alleles having the highest frequencies over all passages and considered the minor allele to be the one with the lowest frequency at the beginning of the experiment (passage 0). We estimated *N* along with locus specific selection coefficients *s*, the sequencing error rate *ϵ* and the per site mutation rate *µ*. We assumed log-uniform priors on *N*, *ϵ* and *µ* such that *log*_10_(*N*) *U*[1,5], *log*_10_(*ϵ*) *= U*[*−*4, *−*0.3] and *log*_10_(*µ*) *= U*[*−*7, *−*1], and a normal prior on the selection coefficients such that *s N*(0,0.05). Since viruses are haploid, we fixed the dominance coefficient at *h =* 0.5. We then run an MCMC using 51 states for 25000 iterations during which each parameter was updated in turn. The first 2000 such iterations were discarded as burn-in phase.

**Simulations** In order to evaluate the power of our method, we simulated data for 20 or 100 unlinked loci with N of 100,1000 or 10000. For each of these settings, we set either 20% or 80% of the loci to be under selection, with an equal representation of four selection coefficients: −0.1, −0.01, 0.01 and 0.1. All loci, both selected and neutral, had the starting allele frequency set at random. The change in allele frequency from one time point to the next was calculated under the Wright Fisher model matching the experimental set-up of our application. Specifically, we simulated a total of 117 generations and took a sample of 1000 sequences every 13 generations.

## Results and Discussion

### Mean Transition Time Approximation

Comparisons of the long term behavior of the here introduced *mean transition time approximation* of the Wright-Fisher process with its discrete realization demonstrate the power of our approximation. In Fig. 2) we show the analytical frequency distribution of alleles with an initial frequency of 0.2 after 10 generations of selection and random drift under the discrete Wright-Fisher process for different population sizes and different selection strength. As expected from our assumptions, the distributions obtained under our approximation have identical means and show only a slightly increased variance for large selection coefficients and a small number of states. A more direct illustration of our assumption is the comparison of the distribution of waiting times for a specific transition. As shown in Fig. 2), our approximation indeed captures the mean transition time perfectly, while again exhibiting an increased variance for large selection coefficients and small number of states.

**Figure 2.**
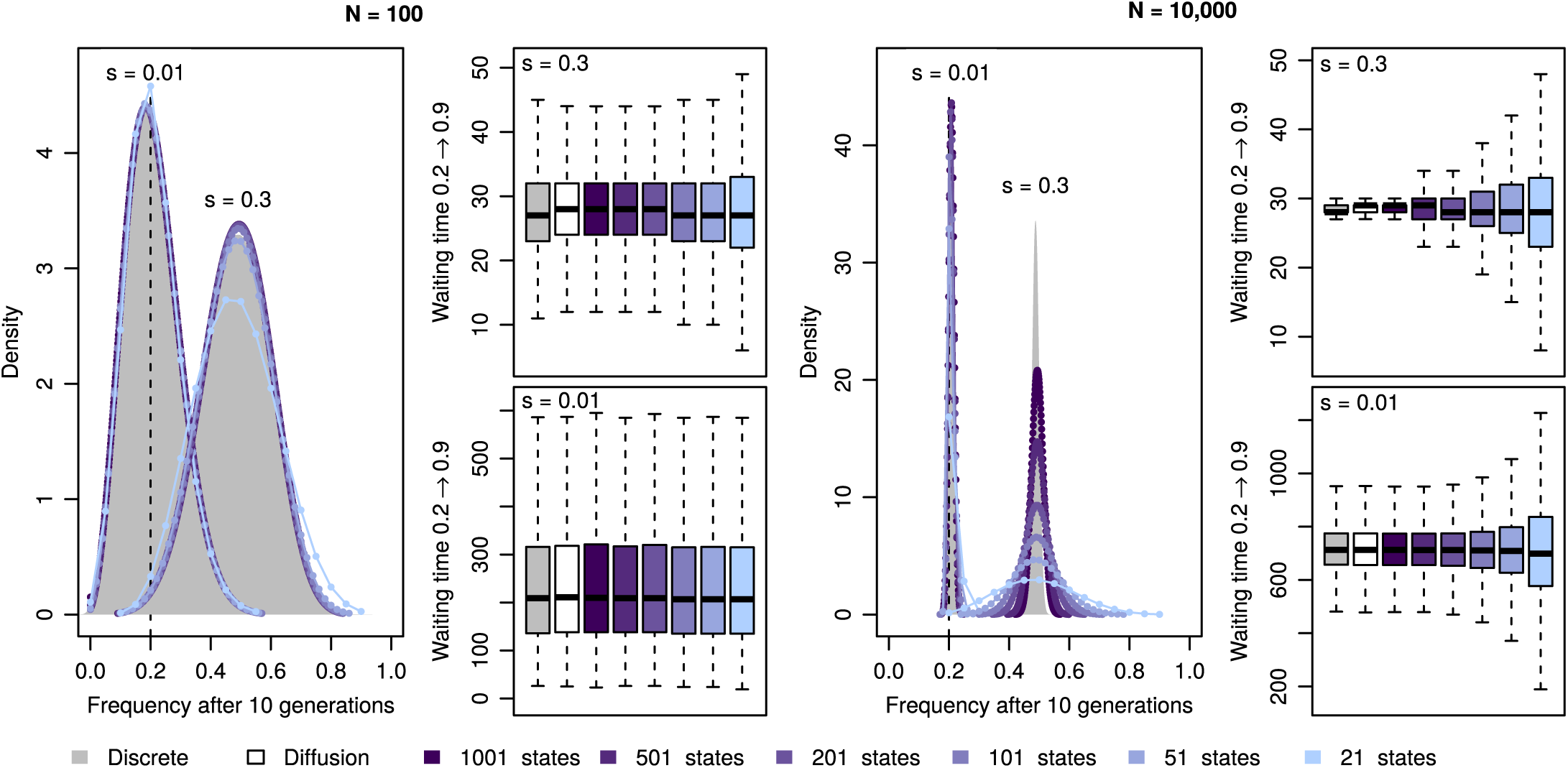
Mean transition time approximation of Markov processes. For both small (N=100, left panel) and large (N=10,000, right panel) population sizes as well as weak (s=0.01) and strong (s=0.3) selection, we show the allele frequency distributions after 10 generations of selection and random drift starting from a frequency of 0.2, as well as the waiting times for a transition from a frequency 0.2 to 0.9. Results obtained under the discrete Wright-Fisher process are given in gray, those obtained under the diffusion approximation considered here in black, and those under our approximation in shades of blue, with darkness increasing with higher number of frequency states considered.

### Power to infer population sizes

While allele trajectories are affected by both selection and drift, we aim here to disentangle these effects by integrating information from multiple loci. We first assessed the power to infer population sizes N accurately under ideal conditions, that is, for 100 unlinked loci in the absence of selection. In Fig. 3 we show the likelihood surfaces for N obtained with different numbers of states, for data simulated under different population sizes. While this analysis suggest high power to infer small population sizes accurately, it highlights the general issue of inferring large population sizes from changes in allele frequencies, accentuated when fewer states are used. The issue arises from the fact that in large populations and over the short time course of evolutionary experiments in general, the changes in allele frequencies between time points are so small, that they are compatible with almost arbitrarily large populations. While using fewer frequency states further decreases the resolution of detectable allele frequency changes, we note that this issue is more general and expected to affect all methods for inferring population sizes from such data, particularly when a small number of samples is used. The best way to overcome this issue is to observe allele frequency changes over larger intervals. Indeed, when taking samples every 130 generations instead of every 13, population sizes up to *N*=100,000 can be estimated accurately (Fig. 3, bottom row).

**Figure 3.**
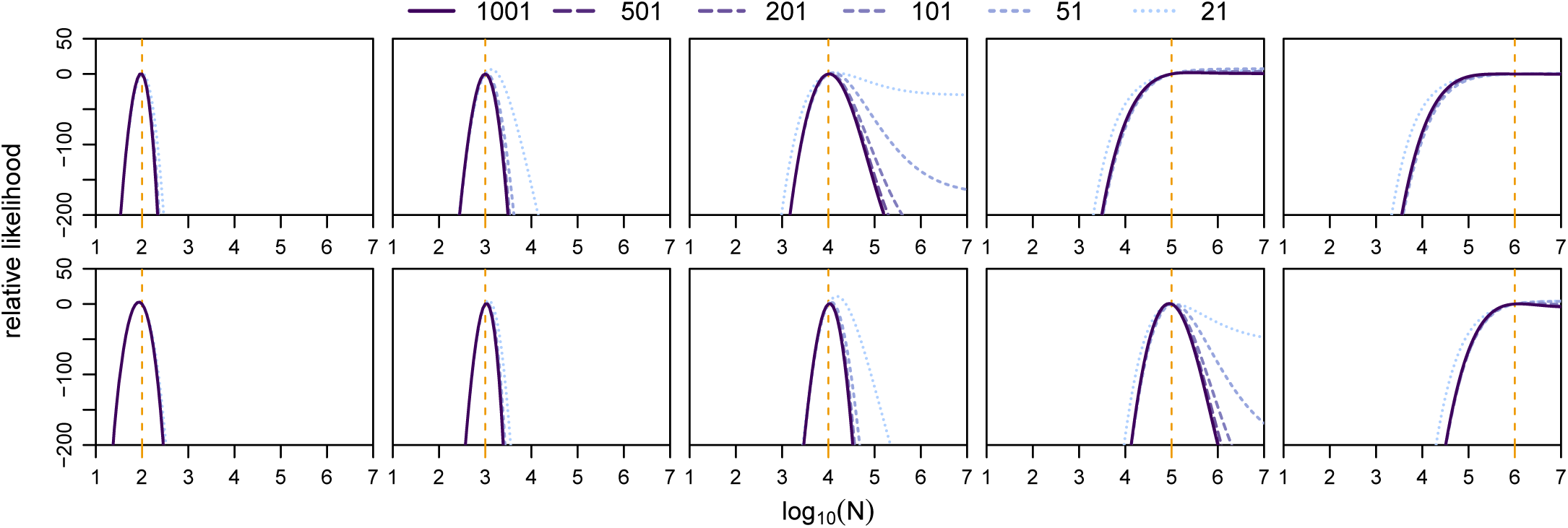
Likelihood surfaces for the population size *N*. Shown are the relative likelihood surfaces obtained via our mean transition time approximation for a particular simulation of 100 neutral loci for different population sizes (vertical dashed lines) and different number of frequency states considered (see legend). The top row is for the case of 13 generations between time points, the bottom row for the case of 130 generations between time points.

### Power to infer selection

To assess the power of our framework to infer locus-specific selection coefficients, we simulated 100 unlinked loci, of which 20% experienced selection at various strengths. As shown in Fig. 4, both the population size as well as the strength of selection affects the power of this inference. For medium to large population sizes, our method infers even small selections coefficients with high accuracy. However, when the population size is small, inference of selection proves more difficult (Fig. 4). While this is generally expected due to the much larger effect of drift in small populations (*Ns* = 10 for the strongly selected alleles), it is accentuated here by our choice to simulate initial frequencies at random. Indeed, when given ideal starting frequencies (0.1 for positively and 0.9 for negatively selected alleles), our method identifies strongly selected alleles accurately even in small populations (Fig. 4).

**Figure 4.**
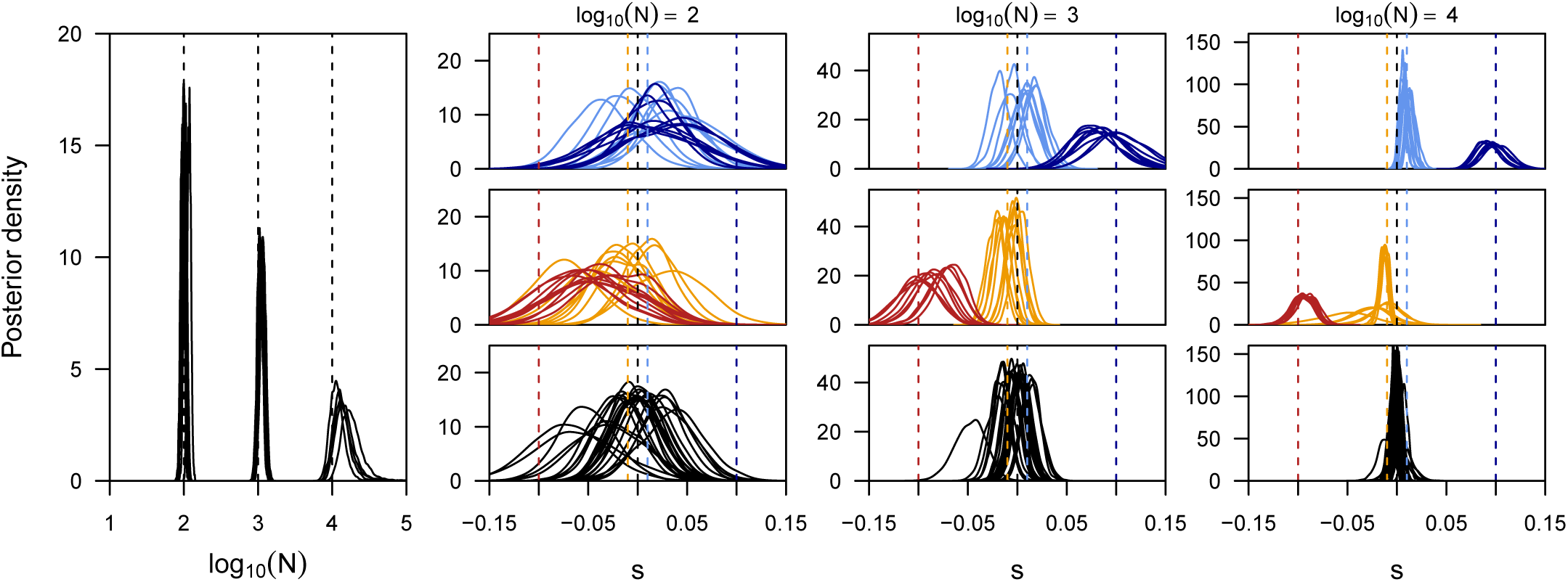
Power to infer selection and demography jointly. Here we show the posterior distributions on the population size (first panel) and locus-specific selection coefficients obtained for five replicate simulations for each of three different population sizes. For each replicate we plot the posteriors of all loci simulated under selection (color) as well as five neutral loci picked at random (black). In all simulations, starting frequencies were chose randomly for each locus.

Remarkably, we found the power to infer the population size as well as locus-specific selection coefficients not to be negatively affected under pervasive selection. This is illustrated by comparing the posterior distributions obtained from simulations where 80% of all loci were targeted by selection (Supplementary Fig. 7) to those shown here where only 20% were affected by selection (Fig. 4). More direct evidence is given in Table 1, where we report the posterior probability for *s >* 0.0 for different combinations of population sizes and selection coefficients and find higher power to identify selected loci under cases of pervasive selection than when only 20% of all loci were simulated under selection.

**Table 1.** Power to identify loci under selection. We report the average and standard deviation (in parenthesis) of the posterior probability 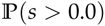 obtained under various population sizes and for the cases of 20% and 80% of all loci simulated under selection.

| fraction selected | $\log_{10}(N)$ | $s=-0.1$ | $s=-0.01$ | $s=0.0$ | $s=0.01$ | $s=0.1$ |
| --- | --- | --- | --- | --- | --- | --- |
| 0.2 | 2 | 0.27 (0.15) | 0.42 (0.24) | 0.50 (0.26) | 0.56 (0.26) | 0.78 (0.18) |
| 0.2 | 3 | 0.00 (0.05) | 0.24 (0.24) | 0.50 (0.30) | 0.74 (0.26) | 1.00 (0.00) |
| 0.2 | 4 | 0.00 (0.00) | 0.06 (0.16) | 0.50 (0.32) | 0.92 (0.17) | 1.00 (0.00) |
| 0.8 | 2 | 0.12 (0.11) | 0.09 (0.14) | 0.50 (0.28) | 0.72 (0.15) | 0.89 (0.15) |
| 0.8 | 3 | 0.00 (0.00) | 0.02 (0.04) | 0.50 (0.38) | 1.00 (0.00) | 0.99 (0.02) |
| 0.8 | 4 | 0.00 (0.00) | 0.00 (0.00) | 0.50 (0.45) | 1.00 (0.00) | 1.00 (0.00) |

### Application to Influenza data

We next applied our approach to publicly available sequencing data of Influenza H1N1 segment 6, obtained at multiple timepoints throughout an evolutionary experiment in which the virus was exposed to an antiviral drug (oseltamivir) (Foll *et al*. 2014a). While allele frequencies are generally estimated with high accuracy due to the very high coverage in this experiment (about 50,000x), sequencing error may contribute substantially to the observed low frequency variants. In addition, many of the observed mutations likely entered the population only during the experiment, but their exact time of origin is blurred by both the sequencing error as well as sampling. We thus extended our framework to estimate the mutation rate as well as the overall sequencing error rate jointly with the demographic and selection parameters.

We applied our extended method to each of the 8 segments of the Influenza genome individually, but obtained highly concordant results among all segments. As shown in Fig. 5, we infer the effective population size during the experiment to be around 7000, a mutation rate of about 10^−5^ and a sequencing error rate of about 10^−3.8^. While our estimates of the mutation and error rates are consistent with published mutation rates for Influenza (Nobusawa and Sato 2006) and RNA viruses in general (Drake *et al*. 1998) and also with the employed quality filters on sequencing reads (Foll *et al*. 2014a), our estimate of the population size is substantially larger than previous estimates of about 225 (Foll *et al*. 2014a). While we found our approach to slightly overestimate larger population sizes under the spacing of time points relevant here, there are several arguments supporting a larger population size. First, the original estimates were obtained under the assumption of neutrality at all loci, while our approach infers *N* jointly with selection. Second, the previous estimates were obtained from a small subset of the data, namely the 147 loci with an observed allele frequency ≤ 1% after down sampling to 1000 reads per locus at no less than three timepoints. In contrast, our inference is based on the raw data at the complete set of 13,395 polymorphic loci, including those with small frequencies particularly informative about drift. Third, the original inference accounted for neither sequencing errors nor mutations. In summary, our results argue for a much larger effective population size than previously reported.

**Figure 5.**
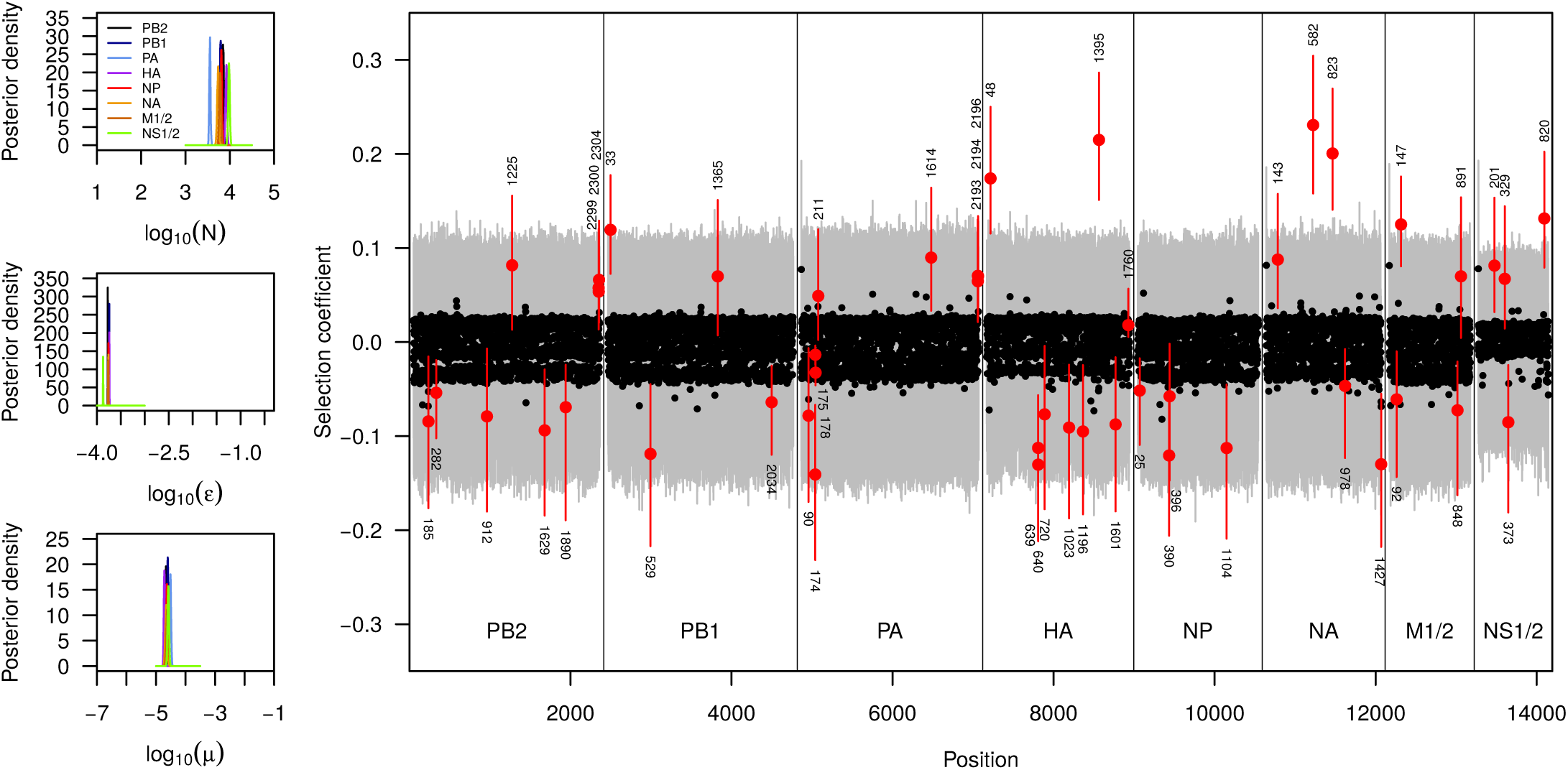
Evolution of drug-resistance in Influenza. Here we show the posterior distributions on the population size (log_10_(*N*)), sequencing error rate (*ϵ*), mutation rate (*µ*) and locus specific selection coefficients *s_l_* estimated independently for each of the six segments of the Influenza genome. For the selection coefficients, black dots represent posterior medians and gray lines indicate the 99% credible intervals. Loci for which the 99% credible interval does not include *s* = 0.0 are shown in red and their actual position within the segment is printed.

Our results on selection, on the other hand, are highly concordant with previous estimates. In Fig. 5 we report the posterior distributions on the locus-specific selection coefficients for all polymorphic sites for each of the 8 segments of the Influenza genome. As expected, most mutations were found to be selectively neutral or under slight purifying selection (observe the slight asymmetry towards negative selection coefficients for many loci). For a few mutations, however, we found compelling evidence for them to be the target of positive selection (99% credible interval does not include 0). On segment NA, there were three such mutations, of which two stand out with an estimated selection coefficient around 0.2. One of these mutations (Y274H) occurred at a locus at which resistance to oseltamivir has been previously described (Collins et ai 2008). Many additional mutations were found to be the target of selection through out the genome, with many of those likely under negative selection. These are mutations that were found at elevated frequencies at the beginning of the experiment, yet at much lower frequencies after a few passages. The complete list of all mutations found to be under selection is given in Supplementary Table 2.

## Conclusion

Here we present a novel, discrete approximation for diffusion processes. This approximation, which we term *mean transition time approximation*, is designed to preserve the long term behavior of the continuous process it approximates, which renders it particularly suitable to study time series data. Here we derived this approximation for the particular case of inferring selection and demography from such time series data under the classic Wright-Fisher model. As shown through extensive simulations, our approximation is well suited to describe allele trajectories through time, even when only a few states are used. This allowed us to develop a Bayesian inference approach to jointly infer the population size and locus-specific selection coefficients with high accuracy. We further extended this model to further estimate the average sequencing error rate, as well as the per generation mutation rate. We finally applied our approach to data from a recent experiment on the evolution of drug resistance in Influenza virus, identifying likely targets of selection and finding evidence for much larger viral population sizes than previously reported.

## Acknowledgments

We thank Nicolas Renzette for his advice on how to identify the protein changes corresponding to individual mutations. This study was supported by Swiss National Foundation grants number PZ00P3_142643 and 31003A_149920 to D.W, and grants from the Swiss National Science Foundation and a European Research Council Starting Grant to JDJ.

## Appendix

## Approximation for large γ = 2Ns

If *γ* = 2*Ns* is large, we get approximations for the Green function which allow for analytic expressions of the integrals. Firstly, we can neglect the minus one terms in the numerator and denominator of (18) and we get the approximation

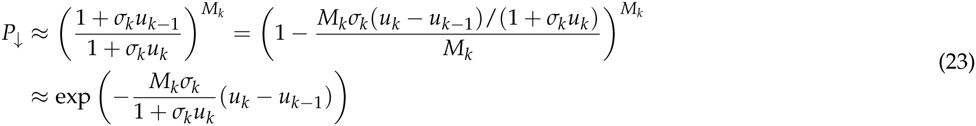
which will be very small for large *γ*. The probability for exit at the upper state is *P*_↑_ ≈ 1. Inserting the first approximating expression or into (19) and using 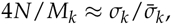, we get

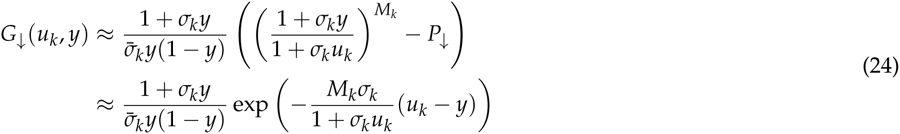

The exponential term is dominant for *y* close to *u_k_*. In the integral we can thus keep the factor of the exponential constant at *y = u_k_* since it does not vary much when *y* is close to *u_k_*:

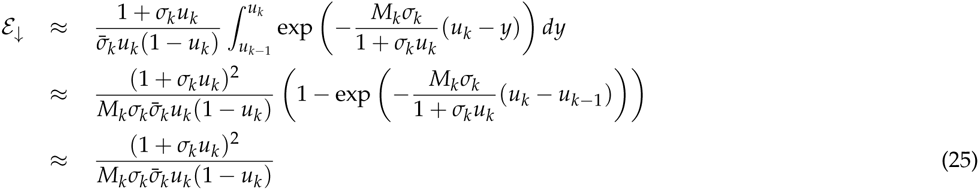

From (20) we get the approximation

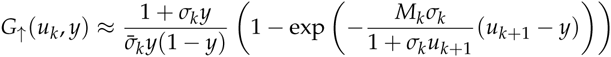

To get *ε*_↑_ we integrate this approximate expression. Observe that the exponential term becomes important only when *y* gets close to *u_k_*+1. For this reason we can safely keep the factor in front of the exponential term constant when integrating the second term:

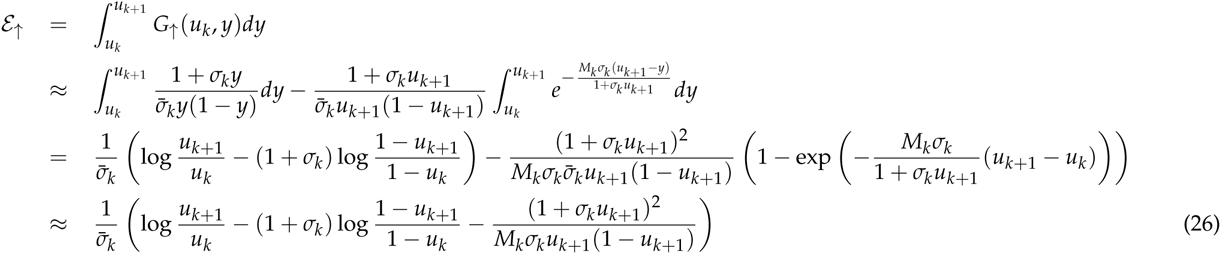

Numerical experiments indicate that the approximative formulae (25) and (26) are adequate when the condition

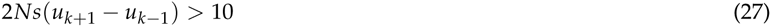
is met. In that case we set *q_k,k−_*_1_ = 0 and

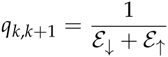

Note that Formula 26 gets singular for *k = K* − 1 since in that case 1 − *u_k_*_+1_ = 0. Using the substitution z = 1 − *y*, we get for that case from (20) the approximation

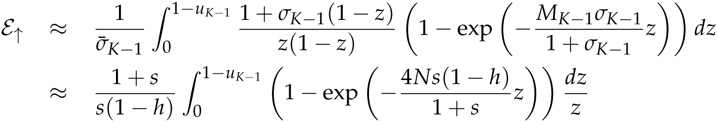

The the last integral can be written as an exponential integral

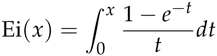
in the form

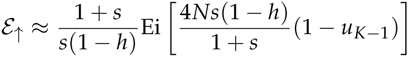

Using the approximation

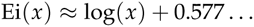
where 0.577… is the Euler-Mascheroni constant, we finally arrive at

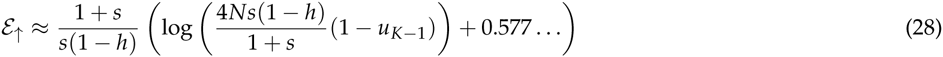

## The Wright-Fisher process in the absence of selection

In the absence of selection (*s* = 0), the expressions for the generator matrix can be explicitly evaluated since *b*(*x*) = 0 (see Eq. 11). We have ϕ(*x*) = *x* and *m*(*y*) = 2*N*/*x*(1 − *x*). From this we get

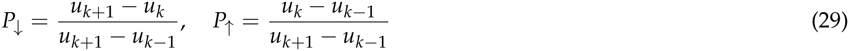

The two parts of the Green function are given by

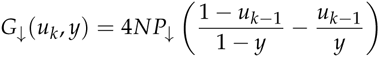
and

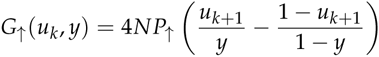

These integrate to

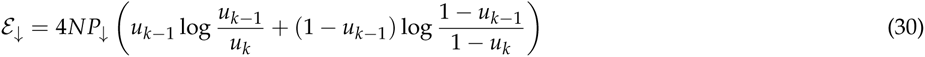
and

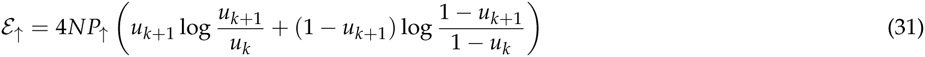

As above we determine the transitions rates by

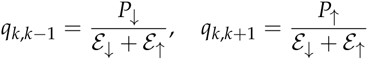

## Supporting Information

**Figure 6.**
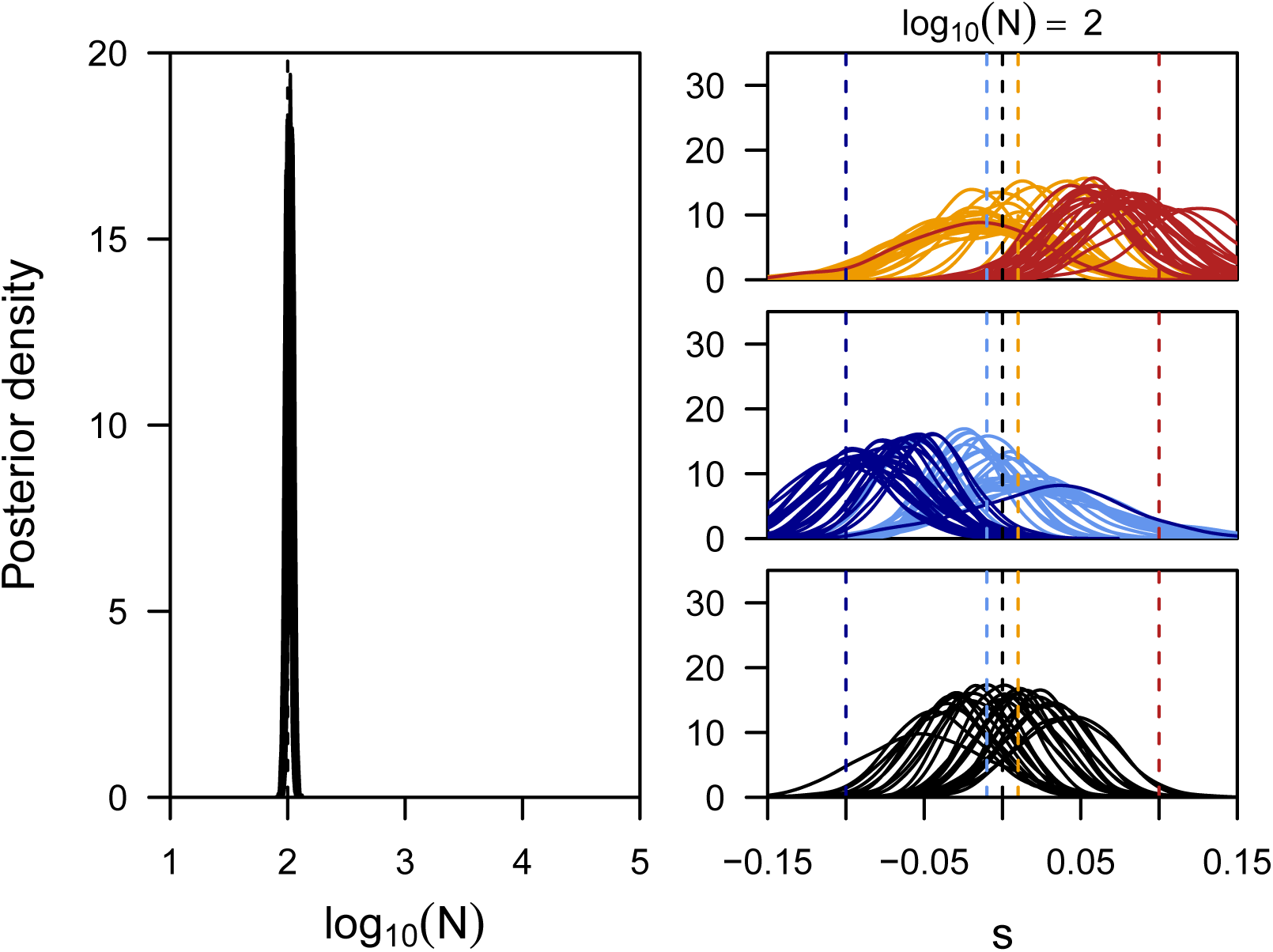
Power to infer selection and demography jointly. Here we show the posterior distributions on the population size (first panel) and locus-specific selection coefficients obtained for five replicate simulations for each of three different population sizes. For each replicate we plot the posteriors of all loci simulated under selection (color) as well as five neutral loci picked at random (black). In contrast to the results shown in the main text, the data was simulated here with more ideal starting frequencies, namely 0.2, 0.5 and 0.8 for positively selected, neutral and negatively selected sites, respectively.

**Figure 7.**
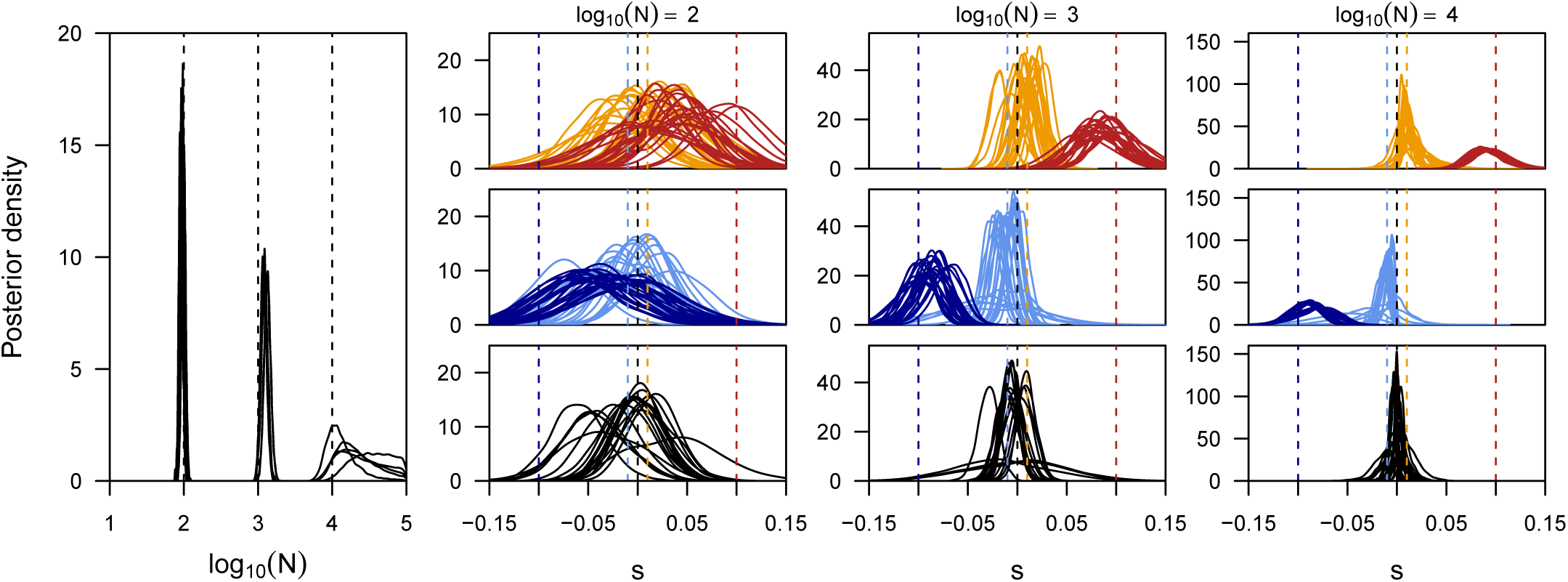
Power to infer selection and demography jointly. Here we show the posterior distributions on the population size (first panel) and locus-specific selection coefficients obtained for five replicate simulations for each of three different population sizes. For each replicate we plot the posteriors of all loci simulated under selection (color) as well as five neutral loci picked at random (black). In all simulations, starting frequencies were chose randomly for each locus. In contrast to the results shown in the main text, 80% of all simulated loci were affected by selection.

**Table 2.** Sites found to be under selection in Influenza

| Segment | Position | Ancestral <sup>a</sup> | Derived | Protein Change<sup>b</sup> | s<sup>c</sup> |
| --- | --- | --- | --- | --- | --- |
| PB2 | 185 | AGG | AAG | R61K | -0.08 (-0.18, -0.02) |
| PB2 | 282 | TCG | TCA | S94 | -0.05 (-0.10, -0.02) |
| PB2 | 912 | GAA | GAG | E304 | -0.08 (-0.18, -0.01) |
| PB2 | 1225 | CGT | AGT | R408S | 0.08 ( 0.01, 0.16) |
| PB2 | 1629 | GAG | GAA | E543 | -0.09 (-0.18, -0.03) |
| PB2 | 1890 | AGA | AGG | R630 | -0.07 (-0.19, -0.02) |
| PB2 | 2299 | - | - | - | 0.06 ( 0.01, 0.12) |
| PB2 | 2300 | - | - | - | 0.05 ( 0.02, 0.11) |
| PB2 | 2304 | - | - | - | 0.07 ( 0.02, 0.13) |
| PB1 | 33 | AAA | AAG | K11 | 0.12 ( 0.07, 0.18) |
| PB1 | 529 | GGT | AGT | G176S | -0.12 (-0.22, -0.04) |
| PB1 | 1365 | AAT | AAC | N455 | 0.07 ( 0.01, 0.15) |
| PB1 | 2034 | AGT | AGC | S678 | -0.06 (-0.12, -0.03) |
| PA | 90 | ACT | ACA | T30 | -0.08 (-0.17, -0.01) |
| PA | 174 | GGT | GGG | G58 | -0.14 (-0.23, -0.07) |
| PA | 178 | CTA | GTA | L59V | -0.03 (-0.05, -0.01) |
| PA | 1614 | GAG | GAA | E538 | 0.09 ( 0.03, 0.16) |
| PA | 2193 | - | - | - | 0.06 ( 0.02, 0.13) |
| PA | 2194 | - | - | - | 0.07 ( 0.04, 0.12) |
| PA | 2196 | - | - | - | 0.07 ( 0.03, 0.13) |
| HA | 48 | CCG | TCG | P6S* | 0.17 ( 0.12, 0.25) |
| HA | 639 | AAT | GAT | N203D | -0.11 (-0.19, -0.06) |
| HA | 640 | AAT | ACT | N203T | -0.13 (-0.21, -0.07) |
| HA | 1023 | GCC | ACC | A331T | -0.09 (-0.19, -0.02) |
| HA | 1196 | ACC | ACT | T388 | -0.10 (-0.18, -0.02) |
| HA | 1395 | AAT | GAT | N455D | 0.21 ( 0.15, 0.29) |
| HA | 1601 | CTA | CTG | L523 | -0.09 (-0.18, -0.02) |
| HA | 1760 | - | - | - | 0.02 ( 0.01, 0.06) |
| NP | 25 | CTC | ATC | L8I | -0.05 (-0.11, -0.02) |
| NP | 390 | ATG | ATA | M130I | -0.12 (-0.21, -0.06) |
| NP | 1104 | AAC | AAT | N368 | -0.11 (-0.21, -0.05) |
| NA | 143 | ACA | ATA | T47I | 0.09 ( 0.04, 0.16) |
| NA | 582 | GGA | GGG | G194* | 0.23 ( 0.16, 0.30) |
| NA | 823 | TAC | CAC | Y274H | 0.20 ( 0.14, 0.27) |
| NA | 978 | TTG | TTC | L326F | -0.05 (-0.12, -0.01) |
| NA | 1427 | - | - | - | -0.13 (-0.22, -0.05) |
| M1/2 | 92 | GAG | TAG | E22stop* | -0.06 (-0.14, -0.01) |
| M1/2 | 147 | GTC | GCC | V41A | 0.13 ( 0.08, 0.18) |
| M1/2 | 848 | TGT | TGG | C274W | -0.07 (-0.16, -0.02) |
| NS1/2 | 201 | AGG | AGA | R67 | 0.08 ( 0.03, 0.15) |
| NS1/2 | 329 | AAA | AGA | K109R | 0.07 ( 0.01, 0.14) |
| NS1/2 | 373 | GAC | AAC | D124N | -0.09 (-0.18, -0.02) |
| NS1/2 | 820 | - | - | - | 0.13 ( 0.08, 0.20) |

<sup>a</sup> Ancestral codon refers to the allele with the highest frequency at the beginning of the experiment (passage 0). Dashes indicate mutations in non-coding regions
<sup>b</sup> Protein changes are reported in standard nomenclature but comparing the derived codon to the ancestral codon (not the published reference).
<sup>c</sup> Reported is the posterior median of the locus-specific selection coefficient, along with the 99% credible interval.

